# BigTop: A Three-Dimensional Virtual Reality tool for GWAS Visualization

**DOI:** 10.1101/650176

**Authors:** Samuel T. Westreich, Maria Nattestad, Christopher Meyer

## Abstract

**Background:** Genome-wide association studies (GWAS) are typically visualized using a two-dimensional Manhattan plot, displaying chromosomal location of SNPs along the x-axis and the negative log-10 of their p-value on the y-axis. This traditional plot provides a broad overview of the results, but offers little opportunity for interaction or expansion of specific regions, and is unable to show additional dimensions of the dataset.

**Results:** We created BigTop, a visualization framework in virtual reality (VR), designed to render a Manhattan plot in three dimensions, wrapping the graph around the user in a simulated cylindrical room. BigTop uses the z-axis to display minor allele frequency of each SNP, allowing for the identification of allelic variants of genes. BigTop also offers additional interactivity, allowing users to select any individual SNP and receive expanded information, including SNP name, exact values, and gene location, if applicable. BigTop is built in JavaScript using the React and A-Frame frameworks, and can be rendered using commercially available VR headsets or in a two-dimensional web browser such as Google Chrome. Data is read into BigTop in JSON format, and can be provided as either JSON or a tab-separated text file.

**Conclusions:** Using additional dimensions and interactivity options offered through VR, we provide a new, interactive, three-dimensional representation of the traditional Manhattan plot for displaying and exploring GWAS data.

## Background

In the last two decades, a decrease in the cost of sequencing has led to a steep increase in the amount of genetic information generated. One aspect of this proliferation of data is an increase in genome-wide association studies (GWAS), each of which requires thousands of individuals to be genotyped or sequenced. In order to interpret the results of a GWAS, it is necessary to condense the large amount of information into a graphic that is still readable and understandable.

The classic visualization for GWAS results is the Manhattan plot (1). Named because of its resemblance to the skyline of a city with a row of tall buildings, the Manhattan plot shows associations for variants across the genome with a given phenotype. Each point displayed on a Manhattan plot represents a single point mutation, or single nucleotide polymorphism (SNP), with the chromosome position plotted along the X axis, and the negative log of the P value for the association test shown on the Y axis. While most measured SNPs have low negative log P values indicating that their associations to the trait being measured by the GWAS are not significant, some SNPs will be highly associated and will thus appear higher on the Y axis (2).

The typical Manhattan plot is useful for providing an overview of the GWAS, showing where significant associations exist on a whole-genome view. These plots are rendered either as static images or as interactive visualizations (3–5). However, a typical two-dimensional Manhattan plot has several drawbacks inherent to its medium: 1) the density of information can potentially obscure interesting results; 2) even in interactive Manhattan plots, selecting a point of interest can be difficult within a dense cluster; 3) additional context such as the population-level allele frequency could aid with interpretation of the results. In addition, standard visualization methods for adding dimensionality (such as varied colors, textures, or shapes) will not work due to the density of information, meaning that adding extra context to a two-dimensional Manhattan plot presents as difficult to impossible.

Of course, traditional static Manhattan plots also lack the ability to zoom in to observe details about specific SNPs, and generally do not provide any identification of individual SNPs unless this information is manually overlaid on the figure through image editing software. Static Manhattan plots also fail to offer additional information about specific SNPs, such as relative abundance or specific chromosomal position. Interactive Manhattan plots offer improvement in many of these areas, but some problems persist due to the natural limitations of two dimensions.

An innovation in technology that is being applied to large genomic datasets is virtual reality (VR). VR applications have been created for several subfields of biology and genetics, including visualization of synteny (6), tracing of neural pathways (7), or three-dimensional protein structure (8–10). These visualizations exist natively in a three-dimensional environment, making them ideal candidates for exploration in virtual reality.

VR is ideal for visualizing large amounts of data that may not be suitable for the constrained display space of two-dimensional monitors. It also permits interaction, allowing for the exploration of data within the figure by observers. A VR-based framework for visualization of genetic or genomic data should be flexible, allowing various datasets to be imported and rendered without requiring any modification of the source code.

One drawback to VR-based visualizations is that creation of the visualization requires a combination of multiple skills. These visualizations are typically created using either WebXR in HTML (6) or an application framework such as Unity or Unreal Engine (8,10), requiring considerable programming experience. Additionally, VR equipment is not yet widely deployed, meaning its availability to researchers may be limited. An ideal VR visualization application should 1) require minimal technical expertise on the user’s part, and 2) be able to display information in a virtual world using a standard monitor.

We created BigTop, a React-based (11) web application that uses the A-Frame framework (12) to render input GWAS summary data in three dimensions. BigTop launches an interactive three-dimensional environment that renders GWAS summary data in three dimensions, wrapping the data in a cylindrical fashion around the user similar to other cylindrical visualizations such as Circos (13). BigTop supports data interaction either through a VR headset or through the combination of a monitor, mouse, and keyboard, allowing users to navigate within the environment and select individual data points to glean more information. Data is read into BigTop in JSON format, but can be provided as a multi-column TSV file and converted to JSON by an included script.

## System Overview

1. GWAS data is provided in JSON format, specifying the chromosome, SNP location on the chromosome, negative log-ten of the p-value, and another measurement used in the z-axis (in all examples, minor allele frequency is used)
  a. For human data, SNP names can be provided. Additionally, a separate preprocessing script can be run on the data to provide information about that SNP’s location (gene name if in a gene). This script only needs to be run once per input file.
  b. For non-human data, a separate file contains chromosome number and size. This can be replaced by the chromosome count and sizes for any other organism.
2. BigTop is easily installed and runs on any system with JavaScript. The display loads in the Chrome or Firefox browsers, and can be viewed through a VR headset with the click of a button. BigTop has been tested and performs on the Oculus Rift, the HTC Vive, and the smartphone-based Google Daydream.
3. BigTop wraps the traditional Manhattan plot around a cylindrical room, placing the user in the center of the room. Chromosomes are marked and colored on the walls, while the height of each point corresponds to the negative log-10 p-value, and the distance along the z-axis (from the center of the room to the wall) indicates the third measurement (minor allele frequency in all example data)[Figure 1].
4. The user can move around the room by taking steps (with a VR headset) or by using the arrow keys (if using a browser). They may control where they look by either moving their head (VR) or by using the mouse to click and drag (browser).
5. In VR, one of the hand controls is set to be a laser pointer (the hand may be switched in BigTop settings). Aiming this laser pointer at a point and pulling the trigger to select that point will display an info panel near the point, providing additional information such as exact p-value, SNP location, gene name, and SNP name (if using human data), and more.
  a. Additionally, the selected point will also extend reference beams to the floor and far wall, better allowing the user to gauge where it falls on the different axes.
6. If using a browser, point selection is possible by centering the point in the center of the user’s vision. A targeting reticule helps align the camera with a point of interest.

## Method/Implementation

### Hardware Requirements

BigTop was tested and developed in non-VR mode on a 2018 MacBook Pro with the following specifications: (1) 3.1 GHz Intel Core i5, (2) 16 GB 2133 MHz LPDDR3 RAM, (3) 512GB SSD, (4) Intel Iris Plus Graphics 650 1536 MB graphics card. BigTop was also tested in VR mode on a laptop computer with the following hardware: (1) Intel i7-7700HQ CPU, (2) 16GB DDR4 RAM, (3) 256GB SSD, (4) GeForce GTX 1060-6GB graphics card. An Oculus Rift was used, with cameras on firmware version 178/e9c7e04064ed1bd7a089, headset on firmware version 709/b1ae4f61ae.

**Figure 1:**
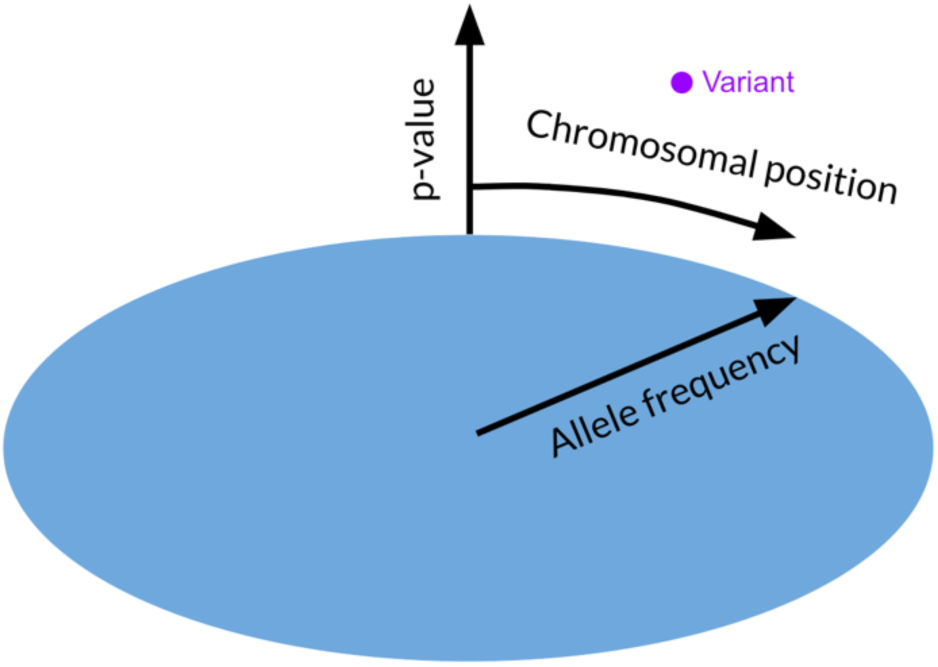
Illustration of the mapping of SNPs in a three-dimensional space. Chromosomal position is indicated by the circumferential location around the edge of the room, while the height of the point indicates the negative log-10 p-value, and the radial distance from the center indicates allele frequency.

### Development Tools

BigTop is coded entirely in JavaScript using the React framework, and makes use of the primitives provided in the A-Frame framework for three-dimensional rendering. A-Frame (12) and other components are installed via the npm package manager. From the Chrome or Firefox web browsers, BigTop may be launched for Oculus Rift, HTC Vive, or another VR display.

### Data Structure & Import

Input data for BigTop is provided in structured JSON format. At minimum, four fields must be provided for each SNP - chromosome number, chromosome position, p-value, and minor allele frequency (MAF). Additional data, such as the SNP name and/or the gene location of the SNP, may also be provided. A Python script provided with BigTop can convert a tab-separated values (TSV) file with these values to JSON format for input into BigTop. Chromosome count and lengths are provided in a chrInfo.json file, and cytoband information is provided in a cytobands.json file. Chromosome count, lengths, and cytoband information is provided for human and rice data.

### Data Rendering & UI

Based on information in the chrInfo and cytobands files, BigTop renders the organism genome as a three-dimensional cylindrical room. Each chromosome is represented by a pie slice of the room, and the cytobands are on each chromosome at approximately eye height. Horizontal and vertical axes are rendered on the floor and on the walls at the intersection between each chromosome.

Input data is used to render each SNP as an object within the VR environment. To improve computational performance, reduce lag, and improve clarity, only SNPs with negative log-10 p-value above a set threshold are rendered as three-dimensional spheres. SNPs with a negative log-10 p-value below the cutoff threshold are instead rendered as circular shadows that are mapped to the floor. The height axis is automatically scaled to reflect the minimum and maximum negative log-10 p-values that are rendered.

Activating a specific SNP by selecting the sphere, either using hand controls in VR or with the cursor, displays additional information about the selected point. A sphere becomes marked as active after being selected in this manner, and an informational panel appears adjacent to the active sphere. This informational panel displays the exact location and p-value of the selected SNP, along with additional information, such as SNP name and gene location, if provided in the input file. Additionally, guides extend from the active sphere to the floor and wall axes to better mark its position in three-dimensional space.

Within the rendered virtual environment, user movement may be performed using arrow keys, if viewing in a browser, or by physically moving while wearing a VR headset.

### Test Datasets

BigTop is capable of displaying GWAS summary data from any organism, as long as the four default values are provided per SNP (chromosome, location, MAF, and p-value). For any organism, chromosome number and sizes must be provided in the chrInfo.json file. BigTop is distributed with two human GWAS datasets; a GIANT dataset which examines SNPs correlated with human height (14), and a breast cancer GWAS from the Nurses’ Health Study (15). To demonstrate its use across multiple species, BigTop also includes a dataset from *Oryza sativa*, examining SNPs related to grain size (16). Toggling between datasets may be accomplished by switching the information in chrInfo.json and cytobands.json, and changing the data file referenced in the main app.js script.

## Results

### Exploring Human & Non-Human Data

The BigTop rendered 3D image allows a user to explore his or her data through interaction; initially loading in the center of the cylindrical image of a GWAS wrapped around the user, the user may move both the location and angle of the camera. As seen in Figure 2, the camera loads facing the start of chromosome 1; in order to view all the data, the user must pivot the camera angle.

**Figure 2:**
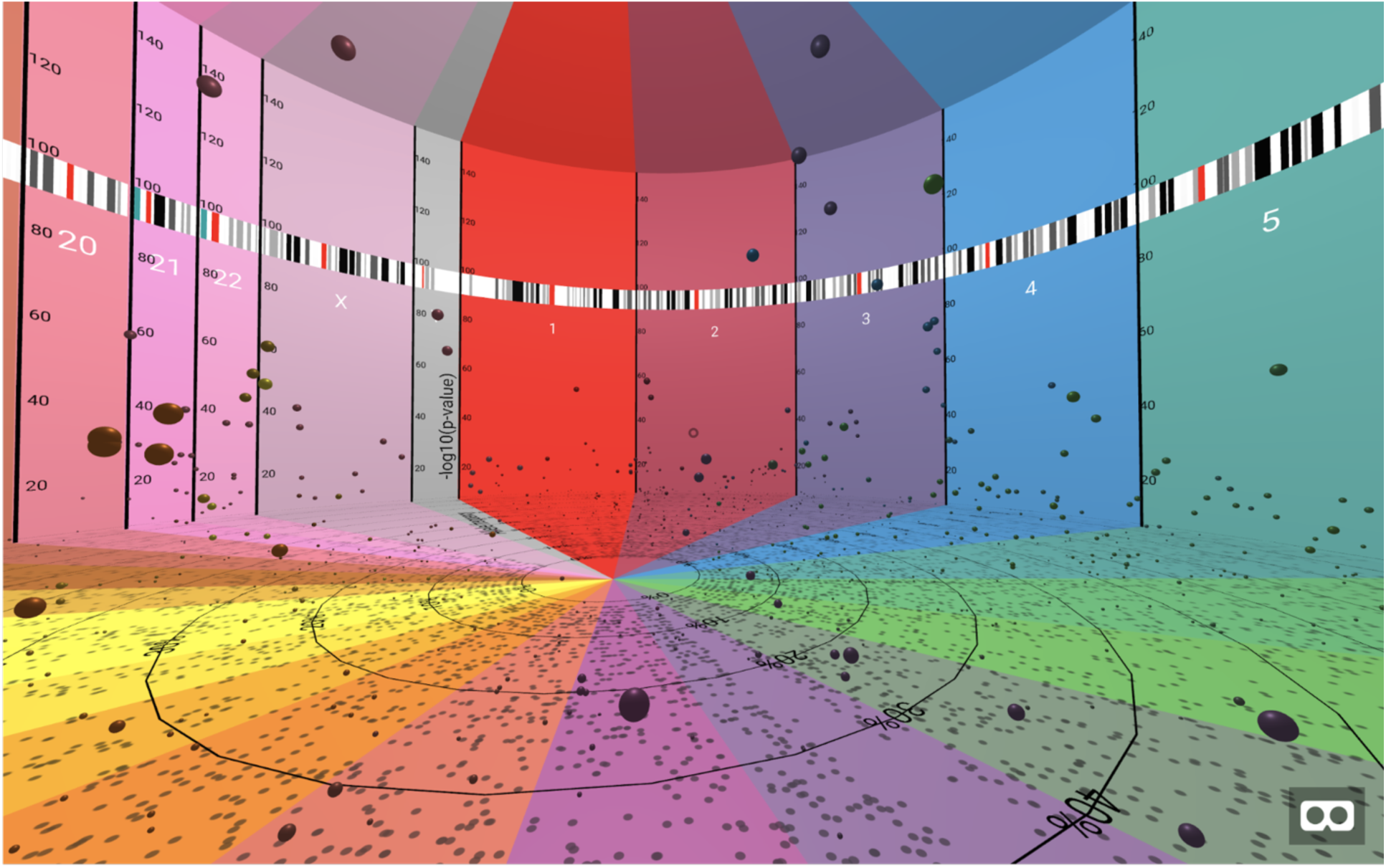
initial view of BigTop upon rendering. The camera loads facing chromosome 1, in the center of the defined “room”. The camera may be rotated to view other areas, either by using the mouse or by turning while wearing an active VR headset.

While the BigTop VR simulation is running, the user may interact with any specific data point by selecting it. If using BigTop with a VR headset and hand controls, a laser pointer is attached to one hand. This laser pointer can be aimed at any individual SNP, represented as a sphere, and an information panel will appear when the trigger is pulled [Figure 3]. If using BigTop with a web browser, a small circle in the center of the screen acts as a heads-up display; centering this circle on any individual sphere will bring up the information panel.

**Figure 3:**
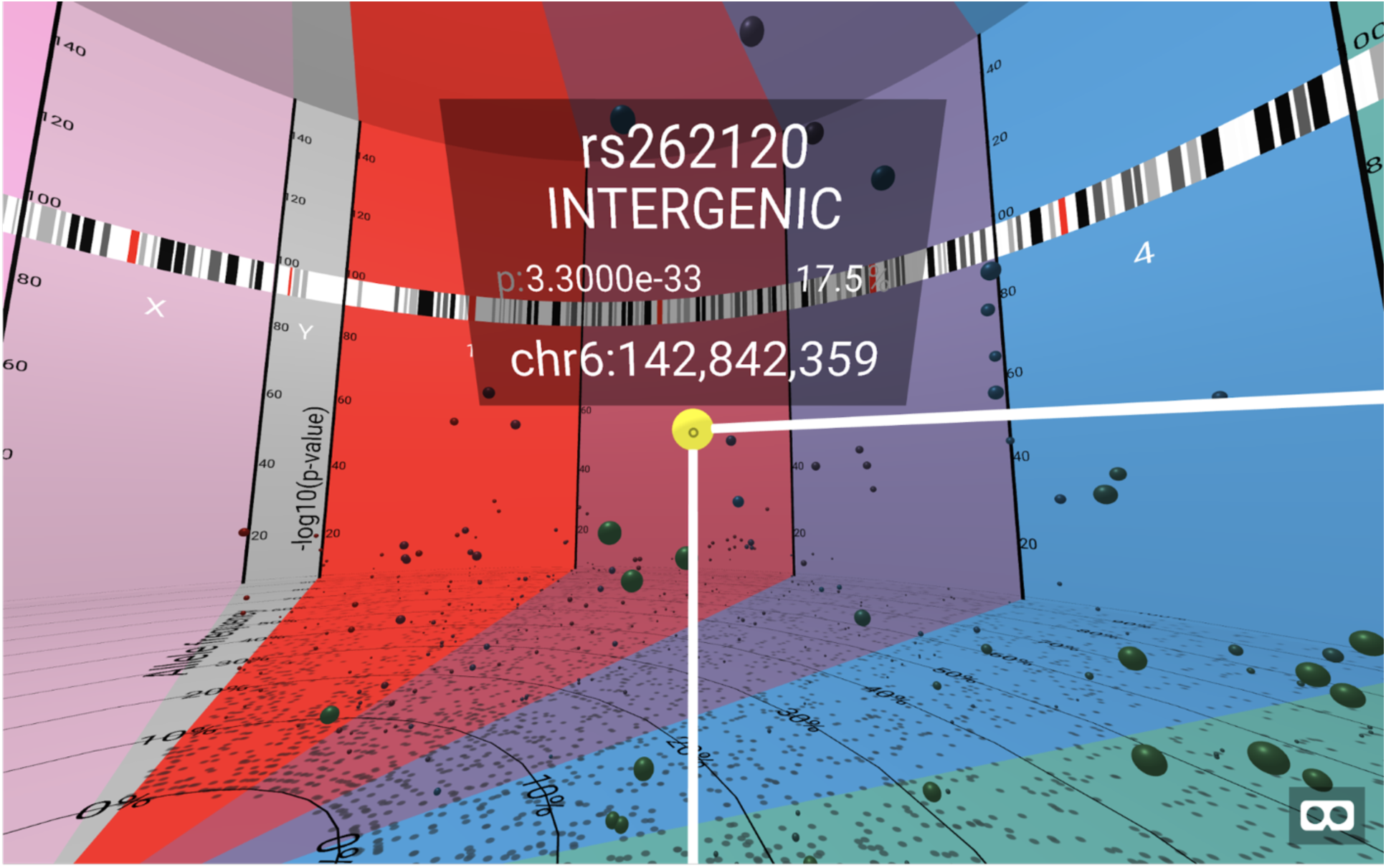
view of an info panel, with a selected SNP. The info panel always displays negative log-10 p-value, chromosome number, and chromosome position, as well as other values (such as SNP name and gene location) if provided in the input file.

Exploration of GWAS data in 3-dimensional space allows for new insights, such as the distribution of SNPs by minor allele frequency. Where a two-dimensional traditional Manhattan plot would show a spike of SNPs at a significant locus, BigTop allows the user to observe the distribution of these SNPs based on their minor allele frequency, or another variable that can be measured and plotted on the Z axis [Figure 4]. This additional dimension of information also allows the user to select specific SNPs for further investigation with the consideration of their relative frequency as an additional weight.

**Figure 4:**
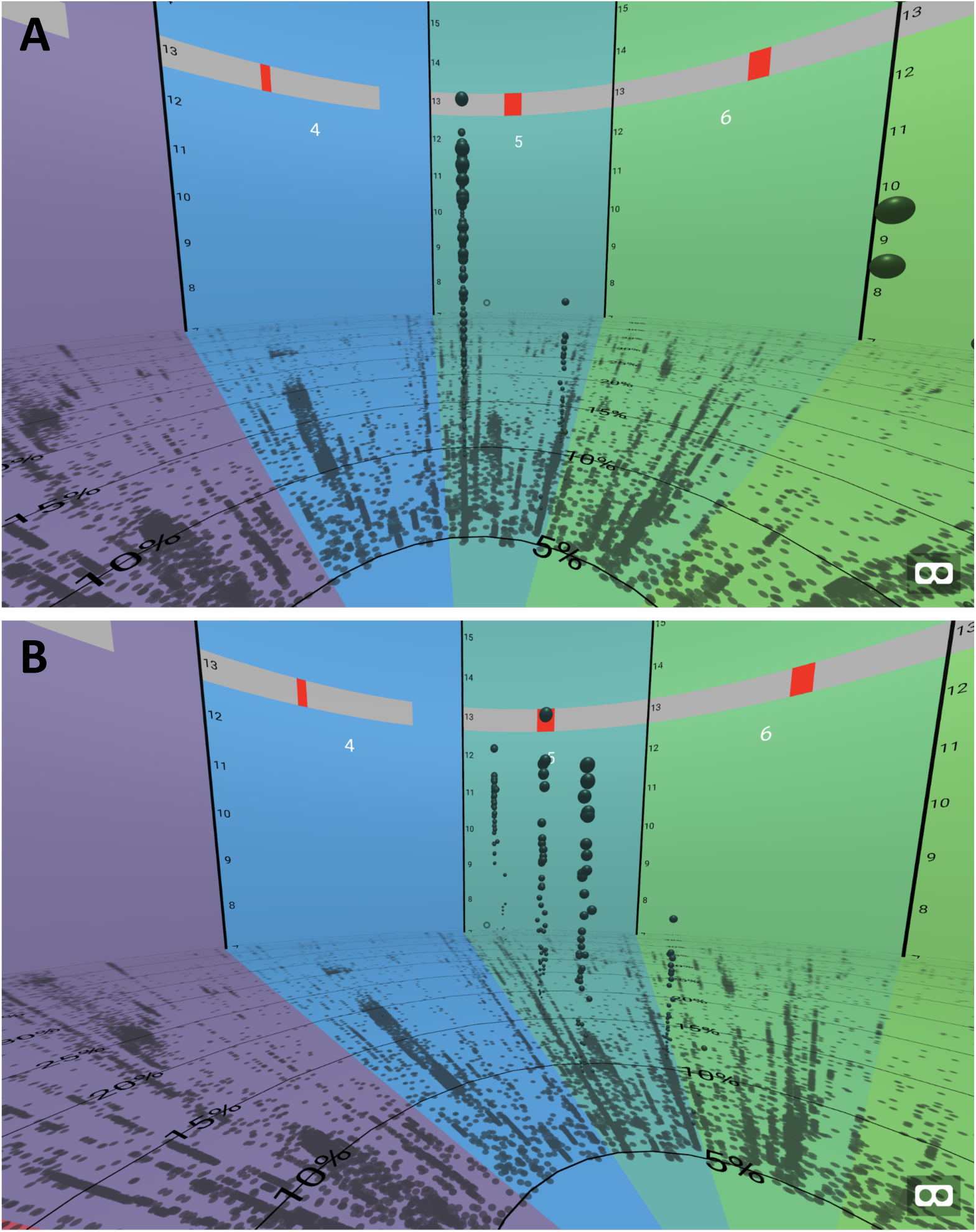
Clusters of significant variants at different frequencies. Multiple variants of significant association were found at clusters of different allele frequencies, suggesting clusters of variants carried together through linkage disequilibrium. This pattern is only visible in three dimensions (right) and not when looking straight down the axis (left) as one would in a regular 2-dimensional Manhattan plot.

### Installation-Free Rendering from a URL

A downside of most VR applications is that they require the user to install and set up an environment on their local machine before he or she may view and explore the VR world. This limitation prevents the VR display from being widely shared or distributed. BigTop circumvents this limitation by allowing users to run the tool by visiting a web page using a browser that supports VR (such as properly-configured installations of Chrome or Firefox). An example of BigTop can be viewed here: https://dnanexus.github.io/bigtop/build/.

This method also allows BigTop results to be easily incorporated into external publications, such as blog posts or scientific papers. The increase in popularity of open access publications and methods has led to an increasing number of researchers making their scripts and methods available on sites such as GitHub. A BigTop visualization can easily be included on such an external site, allowing the researchers to include this interactive figure in their publication.

All configuration in BigTop is done through URL parameters, making it easy to change data sets and customize the user’s experience. Full documentation for getting started can be found at https://github.com/dnanexus/bigtop, but in order to use data sets other than the defaults a user would visit a URL similar to https://dnanexus.github.io/bigtop/build/?data=[dataURL], replacing “[dataURL]” with the URL to the data file to be visualized. (The documentation includes more information on the format of this data file.)

Data files are assumed to be linked to GRCh38, but BigTop will accept any other chromosome and cytoband files. These can also be specified in the URL, such as https://dnanexus.github.io/bigtop/build/?data=[dataURL]&chr=[chrURL]&cyto=[cytoURL], where “[dataURL]”, “[chrURL]”, and “[cytoURL]” all refer to the URLs of the respective files. Again, further information on the formats of these files can be found in the documentation.

## Discussion

Advances in genetic sequencing and analysis technology have led to a greater emphasis on bioinformatics tools for exploring large datasets. Experimental data is now often too large to manually curate and requires specialized software to view and interpret. Presenting high-density visualizations while still providing information in an easily interpreted schema creates additional challenges.

In this paper, we present BigTop, an easy-to-use, interactive VR-based method for exploring genome-wide association study (GWAS) data. This tool aims to increase the information density of the traditional GWAS Manhattan plot, while also allowing interactivity and increased customization and ease of multi-dimensional exploration. BigTop can be run on a local machine or with multiple currently available VR platforms including Oculus Rift, HTC Vive, and Google Daydream. BigTop launches as an HTML object in a Chrome web browser on Windows, Mac, and Unix machines, and can be hosted on an external website, such as GitHub Pages, to allow other users to access and explore the visualization with a link without needing to install or setup any components on their local machine. BigTop reads in a simple comma-separated list, and requires only four components per SNP to render it in three dimensions. BigTop can handle GWAS data from any organism, and settings such as the number and size of chromosomes and pattern of cytoband staining may be altered to suit the chosen organism.

BigTop has several limitations, and there is room for continued improvement and expansion. Currently, BigTop does not support swapping datasets without the need for direct editing of the source code. In a future update, we look to bring a graphical user interface (GUI) to the tool, allowing for the selection of the input dataset, chromosome information, and cytoband pattern information. Additionally, we have further improvements planned, including additional interactivity within the visualization using VR controls.

Advances in the ease and rapidity of gathering GWAS data has made it easier and more affordable than ever before to perform a GWAS to investigate a trait of interest or as part of a larger study. GWAS data is usually visualized in a two-dimensional Manhattan plot, showing the relation between SNP location on the genome and its p-value, indicating how likely it is to be associated with the trait of interest. BigTop allows for visualization of GWAS summary data in three dimensions, including minor allele frequency (MAF) and allowing for users to interactively query any individual point for more information.

## Availability and Requirements

**Project Name:** BigTop

**Project Home Page:** https://github.com/dnanexus/bigtop

**Operating System(s):** Platform independent

**Programming Language:** JavaScript

**Other Requirements:** npm, WebXR-enabled browser (either Google Chrome or Mozilla Firefox)

**License:** MIT license

**Restrictions:** No restrictions

## List of Abbreviations

VR: Virtual reality.
GWAS: Genome-wide association study.
SNP: Single nucleotide polymorphism.
JSON: JavaScript Object Notation.
MAF: Minor allele frequency.
TSV: Tab-separated values.

## Declarations

### Ethics approval and consent to participate

Not applicable.

### Consent for publication

Not applicable.

### Availability of data and material

All components of BigTop are available publicly on GitHub at https://github.com/dnanexus/bigtop. The repository includes documentation and instructions for installation and running the program. The repository additionally includes public data that can be rendered.

All starting data used in development and implementation of BigTop are publicly available. Summary datasets used by BigTop for visualization are available with the source code in the public repository.

### Authors’ contributions

Initial design of BigTop provided by SW, MN, and CM. Code was written and developed by CM and MN. Data was obtained and structured by SW. All authors contributed to and approved the final manuscript.

## Acknowledgements

The authors wish to thank their colleagues at DNAnexus who aided in the development and testing of the BigTop framework, including Andrew Carroll, who provided funds for the purchase of testing equipment.

